# Iota-carrageenan protects the ocular surface from desiccation-induced cell death and tissue damage in vitro and ex vivo

**DOI:** 10.64898/2026.03.10.709798

**Authors:** Sabine Roch-Nakowitsch, Antonella Russo, Christiane Koller, Andrea Dolischka, Marielle König-Schuster, Hanna Dellago, Eva Prieschl-Grassauer

## Abstract

Iota-carrageenan is a natural polymer with moisturizing, mucoadhesive and shear-thinning properties. In this study, we aimed to evaluate the protective effects of iota-carrageenan on ocular surface against dehydration, to demonstrate its suitability for the use in lubricant eye drops. We utilized a human epithelial corneal cell culture model to test if pre-incubation with iota-carrageenan solution could protect cells from desiccation-induced cell death, to compare its effect with other natural polymers commonly used in artificial tear products, and to determine the optimal iota-carrageenan concentration. An *ex vivo* porcine eye model was established to confirm the protective effect of iota-carrageenan against dehydration on ocular tissue. Pre-incubation with 1.2 mg/ml iota-carrageenan increased the survival half-life of human corneal epithelial cells upon dehydration by three-fold; the effect was in the same range as observed for large molecular weight hyaluronic acid, and superior to all other tested natural polymers. The highest tested concentration of iota-carrageenan, 1.6 mg/ml, extended the cellular survival half-life by eight-fold while maintaining healthy cellular morphology. Repeated *ex vivo* instillation of an iota-carrageenan-based ophthalmic formulation into porcine eyes significantly protected the ocular surface from desiccation-induced corneal damage, as shown by corneal fluorescein staining These data suggest that Iota-carrageenan effectively moisturizes and protects the ocular surface, supporting its potential as a promising novel ingredient for eye drops in the management of dry eye disease.

## 2. Introduction

Dry eye disease (DED), also termed dry eye syndrome (DES) or keratoconjunctivitis sicca (KCS), is a multifactorial disease of the ocular surface characterized by a loss of homeostasis of the tear film and tear film instability. Symptoms include ocular dryness, irritation, redness, discharge, and blurred vision, can range from mild and occasional to severe and continuous, and have been shown to negatively impact work productivity and daily activities (Kanellopoulos and Asimellis, 2016; Liu et al., 2022; Sheppard et al., 2023).

The tear film of the eye, consisting of the lipid layer, the aqueous layer, and the mucous layer, is responsible for the maintenance of the corneal epithelium. The aqueous layer is responsible for the osmolality of the ocular surface and changes thereto lead to a disturbed homeostasis of the tear film. The mucous layer has viscoelastic and lubricating properties and detains the moisture on the corneal epithelium and is responsible to form a barrier against drugs, toxins, virus particles or allergens (Račić and Krajišnik, 2023; Willcox et al., 2017).

Mild to moderate cases of DED are usually treated with lubricant eye drops, gels or ointments that can be easily purchased over the counter. The majority of the available OTC lubricating products are gel-like ophthalmic formulations composed of ingredients that contain multifunctional large biopolymers such as hyaluronic acid, carboxymethyl cellulose or hydroxypropylmethylcellulose in order to increase the viscosity of the tear film, to moisturize and prolong the residence time of an aqueous layer on the cornea (Cassano et al., 2021; Račić and Krajišnik, 2023; Zhang et al., 2017). An ideal compound for the treatment of DED that aims to enhance tear film stability should possess properties similar to the mucous layer and in addition should be mucoadhesive to remain on the ocular surface. Here, we propose iota-carrageenan as a promising, safe and effective new alternative for the treatment of DED. Carrageenans are natural polysaccharides built of D-galactose and 3, 6-anhydro-D-galactose units which are extracted from certain red seaweeds of the Rhodophyceae class. In the food, cosmetic and pharmaceutical industry, carrageenans are widely utilized due to their excellent physical functional properties, such as gelling, thickening, emulsifying and stabilizing abilities (Li et al., 2014). There are three main types of carrageenan based on the number of sulfate groups attached to the galactose molecule, which differ with regard to their solubility and their gelling and thickening capacity: kappa- (one sulfate group per two galactose molecules), iota- (two sulfate groups per two galactose molecules) and lambda- (three sulfate groups per two galactose molecules) carrageenan. The toxicological profile of carrageenans has been thoroughly evaluated, and they have been established to have minimal or no adverse physiological effects (Cohen and Ito, 2002). No health hazards have been documented for iota-carrageenan so far, and it has been designated as “Generally Recognized As Safe” (GRAS status) by the FDA for food and topical applications (U.S. Food & Drug Administration, SCOGS (Select Committee on GRAS Substances), 2020).

Iota-carrageenan is certified for marketing in the EU, parts of Asia and Australia under the brand name Carragelose® as a component of nasal sprays, throat sprays and lozenges. Previous pre-clinical and clinical studies led by us and others have demonstrated the broad, non-specific antiviral and barrier-forming properties of iota-carrageenan (Bichiri et al., 2021; Eccles et al., 2010; Fazekas et al., 2012; Fröba et al., 2021; Graf et al., 2018; Grassauer et al., 2008; Koenighofer et al., 2014; Ludwig et al., 2013; Morokutti-Kurz et al., 2021; Unger-Manhart et al., 2024).

In addition, iota-carrageenan has great potential to be used in lubricating and moisturizing eye drops based on the following properties: Firstly, it forms an *in situ* gelling system upon exposure to physiological conditions (Akash Muktiram Shendge*, Vikas B Wamane, Jayshree R. Shejul, 2024; Rupenthal et al., 2011) and hence retains moisture on the ocular surface due to its excellent water-binding capacity (Thommes et al., 2009), counteracting evaporation. Secondly, it is a viscosity enhancer that exhibits viscoelastic behavior (Hilliou, 2021; HOSSAIN et al., 1997). Thirdly, its mucoadhesive properties are similar like that of the mucous layer (Račić and Krajišnik, 2023; Vigani et al., 2019; Volod’ko et al., 2024). The second and third property additionally improves the retention time on the ocular surface and thereby may provide a prolonged stay on the ocular surface compared to aqueous solutions. Fourthly, it is a polymer with non-Newtonian properties and as such it is per definition shear-thinning, hence reducing resistance against movements of the eyelid when blinking (Ludwig, 2005). Finally, iota-carrageenan acts locally and is not absorbed by the body, therefore it may be used also during pregnancy and lactation.

The suitability of iota-carrageenan for ophthalmic applications has already been demonstrated in various contexts: Carrageenans have been tested and proposed as promising carriers for ocular drug delivery (Bonferoni et al., 2004; Omran et al., 2023; Volod’ko et al., 2024). A commercial iota-carrageenan-containing viscous gel has been proven to be safe and effective when used to protect and hydrate the corneal surface during ophthalmic surgery (Giardini et al., 2018; Mencucci et al., 2019). Hence, the potential of iota-carrageenan in multiple ophthalmic therapeutic areas may deserve further attention. But to date, no study has evaluated the suitability of iota-carrageenan as lubricant and moisturizing compound for treatment of DED.

The present research evaluated the moisturizing properties and the protective effects of iota-carrageenan against air-induced desiccation. Using both *in vitro* cell culture and *ex vivo* porcine eye model systems, we assessed the protective effect of iota-carrageenan in comparison to established substances commonly used for the treatment of DED. Additionally, we demonstrated a concentration-dependent protective effect of iota-carrageenan against dehydration and determined the concentration of iota-carrageenan for use in an ophthalmic formulation.

## 3. Methods

### 3.1 Cell culture

We have adapted a cell culture model that has previously been used in the development of DED treatments to assess prevention of cell death from desiccation (Hill-Bator et al., 2014). Immortalized Human Corneal Epithelial Cell Line (IM-HCEpiC; Innoprot, Derio (Bizkaia), Spain; # P10871-IM) were cultured according to the supplier’s instructions. Briefly, cells were maintained in IM-Ocular Epithelial Cell Medium (Innoprot; P60189) supplemented with growth factors and antibiotics in standard tissue culture flasks. Cells were incubated at 37°C in a humidified atmosphere containing 5% CO_2_. The culture medium was exchanged with fresh medium every 2-3 days, and cells were passaged upon reaching 80-90% confluency using trypsin-EDTA.

### 3.2 *In vitro* dehydration assay

Human immortalized cornea epithelial cells (IM-HCEpiC) were seeded into 96-well plates (TPP; 92696) at a density of 1x10^4^ cells/well on the day before the experiment. The next day, medium was replaced with the sample solutions described below or with fresh cell culture medium only as control, and incubation under cell culture conditions was continued for 2 h. Subsequently, supernatants were removed, and culture plates were placed without lid in a dry incubator at 33°C for defined timespans of 2-40 min to allow dehydration. The following controls were included in the experiment: i) internal positive control (iPC) – cells were incubated with medium but not submitted to dehydration; ii) sample positive control (sPC) – cells were incubated with respective sample solution but not submitted to dehydration; iii) negative control (NC) – cells were incubated with cell culture medium and submitted to dehydration. Cell viability following dehydration was assessed fluorometrically using a resazurin based assay (Alamar Blue, Acros; 189900010). Cells were incubated with a 1x Alamar blue solution for 30 min at 37°C in a humidified atmosphere containing 5% CO_2_. After incubation, the fluorescence of reduced resazurin was measured using a fluorescence plate reader (BMG Fluostar Omega) with excitation at 544 nm and emission at 590 nm. Cell viability was calculated for each sample relative to the corresponding sPC viability. For each tested sample solution, a dose-response analysis was performed and the respective cellular survival half-life, i.e., the time point when 50 % of the metabolic signal was still detectable, was calculated based on the dose-response curves. The goodness-of-fit of the dose-response analysis is expressed as confidence interval of the calculated half-life.

### 3.3 Sample solutions used in the in vitro dehydration assay

For comparison of the dehydration-protective effect of natural polymers, the following substances were dissolved in distilled water (aqua destillata, AD; Aqua Braun; B.Braun; 0082479E) and then mixed 1:1 with cell culture medium to achieve a final assay concentration of 1.2 mg/ml: fucoidan (two types: derived from Fucus vesiculosus (Sigma; F8190) and from Undaria pinnatifida (Sigma; F8315) ), dextran sulfate sodium salt (Sigma; D6001), carboxymethyl cellulose (Hercules; 7MF Pharm), hydroxypropyl methylcellulose (Fagron; 102517), hyaluronic acid (three different molecular weight species: 300-500 kDa (Sigma; 00941), 750-1000 kDa (Sigma; 53163) and 1.2-1.5MDa (Sigma; 53747), kappa-carrageenan (Du PontNutrition), and iota-carrageenan (Du PontNutrition).

For investigation of an iota-carrageenan dose effect, iota-carrageenan was dissolved in AD at a concentration of 3.2 mg/ml and this stock solution was further diluted with AD to prepare solutions with concentrations down to 0.15 mg/ml. These iota-carrageenan solutions were mixed 1:1 with cell culture medium to obtain the following final assay concentrations of iota-carrageenan: 1.6, 1.4, 1.2, 1.0, 0.8, 0.6, 0.3, 0.15, and 0.075 mg/ml.

### 3.4 Cytoskeletal and nuclear staining after treatment and dehydration

Cell treatment and dehydration: IM-HCEpiC were seeded at a density of 8x10^4^ cells/well on coverslips (VWR; ECN631-1578) in a 24-well plate (TPP; 92024) the day before the experiment. Cells were incubated for 2 h with 3.2 mg/ml iota-carrageenan in an isotonic buffer, mixed 1:1 with cell culture medium to give a final assay concentration of 1.6 mg/ml. After the 2 h incubation, specific wells were designated for dehydration condition. Cell culture medium and treatment solution was aspired from designated wells. Subsequently, the plate was placed without lid in a dry incubator at 33°C for 10 or 30 min. Following dehydration, cells were washed with PBS (Sigma; D8537), fixed with Roti Histofix (Roth; P087.6) for 10 min, washed with PBS (3x), permeabilized with 0.1% Triton X-100 (Serva Feinbiochemica; 37240) for 5 min, and washed with PBS (3x). Coverslips were then stained for 1 h with 1x Phalloidin-iFluor 488 (Abcam; AB176753) plus 1 µg/ml of DAPI (Sigma; D9542) in PBS plus 1% BSA (Roth; 8076.4) on parafilm in a humidified petri dish. Coverslips were washed with PBS (3x) and mounted on glass slides (Henry Schein; 900-2573) with mounting medium (Roti-Mount Aqua; Roth; 2848.1) or epifluorescence microscopy (Axio ExaminerD1 with axiocam 202 camera; Carl Zeiss).

### 3.5 *Ex vivo* dehydration assay

The induction of ocular surface damage by desiccation is based on a published model (Choy et al., 2004). Porcine eyes including the surrounding tissue of the eye socket were extracted from skulls of pigs (German-Large-White) slaughtered at a local slaughterhouse. Eyes were examined for corneal damage by distributing one drop (25 µL) of sodium fluorescein (Fluka, 46960; 0.1% w/v in PBS) on the ocular surface and scoring the surface damage under a black light torch l λ _Em_: 395 nm according to the Oxford grading scale. Only undamaged eyes (Oxford scale grading 0 or 1) were used for the experiment. Eyes judged as undamaged were pinned open and incubated in a laminar flow hood for 6 h at room temperature and approx. 50% humidity. Every 15 min during this 6 h period, a single drop of sample solution (described below) or 0.9% NaCl (B.Braun, 3570130) (negative control), was dropped onto the cornea and spread across the surface by blinking with the nictitating membrane.

At the end of the 6 h dehydration period, eyes were again submitted to Oxford grading after instillation of 1 drop of sodium fluorescein (0.1% w/v in PBS). Next, eyeballs were excised from the surrounding tissue and immersed into sodium fluorescein (0.1% w/v in PBS) for 3 min, followed by ten rinsing steps in PBS. Corneas were dissected, and 0.8 cm diameter punches were taken from the center. Fluorescein was extracted from cornea punches in Triton X-100 (Sigma, T8787-50mL) (1 % v/v in PBS) at 4°C for 48 h. Relative fluorescence units of extracts were measured at Ex/Em 485/520 nm using a BMG Fluostar Omega plate reader, and fluorescein concentration [µg/g] of each sample was calculated based on a 20-point standard curve with two-fold serial dilutions. P values were derived from one-way ANOVA with Tukey’s post-hoc test.

### 3.6 Sample solutions used in the *ex vivo* dehydration assay

Iota-carrageenan dissolved in isotonic buffer composed of disodium-EDTA (Titriplex III) 1.00 mg/ml, citric acid monohydrate 0.46 mg/ml, disodium hydrogen phosphate dihydrate 3.67 mg/ml, sorbitol 40.00 mg/ml, at concentrations 0.6 mg/ml, 2.0 mg/ml, and 3.2 mg/ml.

## 4. Results

Based on the inherent properties of iota-carrageenan and its demonstrated suitability for ophthalmic applications [Bonferoni et al., 2004; Giardini et al., 2018; Mencucci et al., 2019; Omran et al., 2023; Volod’ko et al., 2024], we set out to examine the potential of iota-carrageenan for use in the treatment of DED.

To quantify and contextualize the moisturizing capacity of iota-carrageenan, we employed a cell culture dehydration assay using human immortalized cornea epithelial cells (IM-HCEpiC). Cells were-preincubated for 2 h with iota-carrageenan or other polymers that are currently used in products for the treatment of DED, all at a final assay concentration of 1.2 mg/ml. Subsequently, cells were submitted to dehydration for various time periods, and viability after dehydration was determined fluorometrically using a resazurin based assay. Cellular survival half-lives upon dehydration, i.e. the duration of dehydration when cell viability reached 50% of the respective non-dehydrated control, were calculated for each sample. As shown in **figure 1**, pre-incubation of IM-HCEpiC with iota-carrageenan protected cells from desiccation-induced cell death and led to a threefold increase of the cellular survival half-life upon dehydration. This effect was specific to iota-carrageenan and was not observed with kappa-carrageenan. The magnitude of effect was comparable with that of large molecular weight hyaluronic acid (1.2-1.5 MDa) and was superior to the effect of all other tested polymers, including fucoidan, dextran sulfate, carboxymethyl cellulose, hydroxypropyl methylcellulose, as well as low and medium molecular weight hyaluronic acid (0.3-1 MDa).

**Figure 1:**
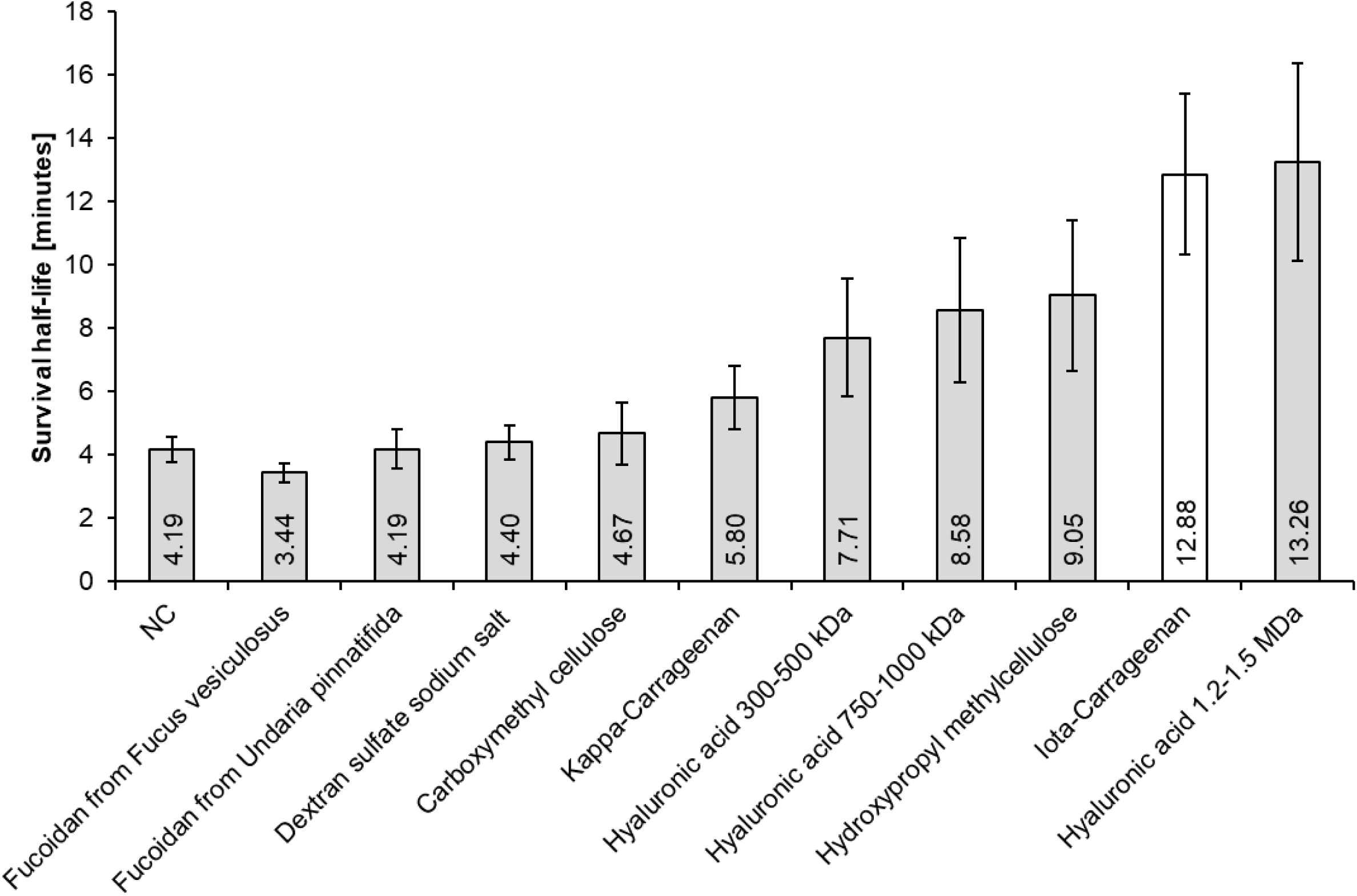
Iota-carrageenan (white column) protect corneal cells from dehydration. IM-HCEpiC were pre-incubated with natural polymers in cell culture medium at a concentration of 1.2 mg/ml for 2h and then dehydrated for 2-40 min by removing the medium and placing the culture plates without lid in a dry incubator. Viability was assessed after dehydration photometrically using a resazurin based assay. Survival half-life was defined as time after which 50% of cells were still metabolic active. Error bars represent 95% CI (n= min. 3). NC = negative control, cells pre-treated with cell culture medium w/o polymers, followed by dehydration.

Next, we aimed at determining if there is a concentration dependence of iota-carrageenan for protection against dehydration, again using the IM-HCEpiC *in vitro* dehydration model. In this case, cells were-preincubated for 2 h with various concentrations of iota-carrageenan spanning final assay concentrations from 1.6 mg /ml down to 0.075 mg/ml, followed by dehydration for various durations, and viability assessment based on resazurin (Alamar Blue). Our results demonstrate that pre-incubation with iota-carrageenan enhanced cell viability after dehydration in a dose-dependent manner. As shown for selected concentrations in **Figure 2a**, cells pre-incubated with medium alone could compensate for only a 2 min dehydration period without loss of viability and had already lost 50% viability within less than 5 min of dehydration. In contrast, cells pre-incubated with the highest tested concentration of 1.6 mg /ml iota-carrageenan were able to withstand dehydration and maintain high viability levels of > 90% for at least 20 min and showed an increase in half-life survival by eight-fold in comparison to the negative control (**Figure 2b**). Lower iota-carrageenan concentrations had similar, though proportionally weaker effects. Iota-carrageenan concentrations below 1 mg/ml led to markedly reduced viability already after 5-10 min of dehydration but still extended the dehydration survival half-life, albeit to a lower extent. Concentrations below 0.15 mg/ml yielded similar results as the negative control (data not shown). In sum, the protective effect of iota-carrageenan against dehydration shows a clear dose-dependence.

**Figure 2:**
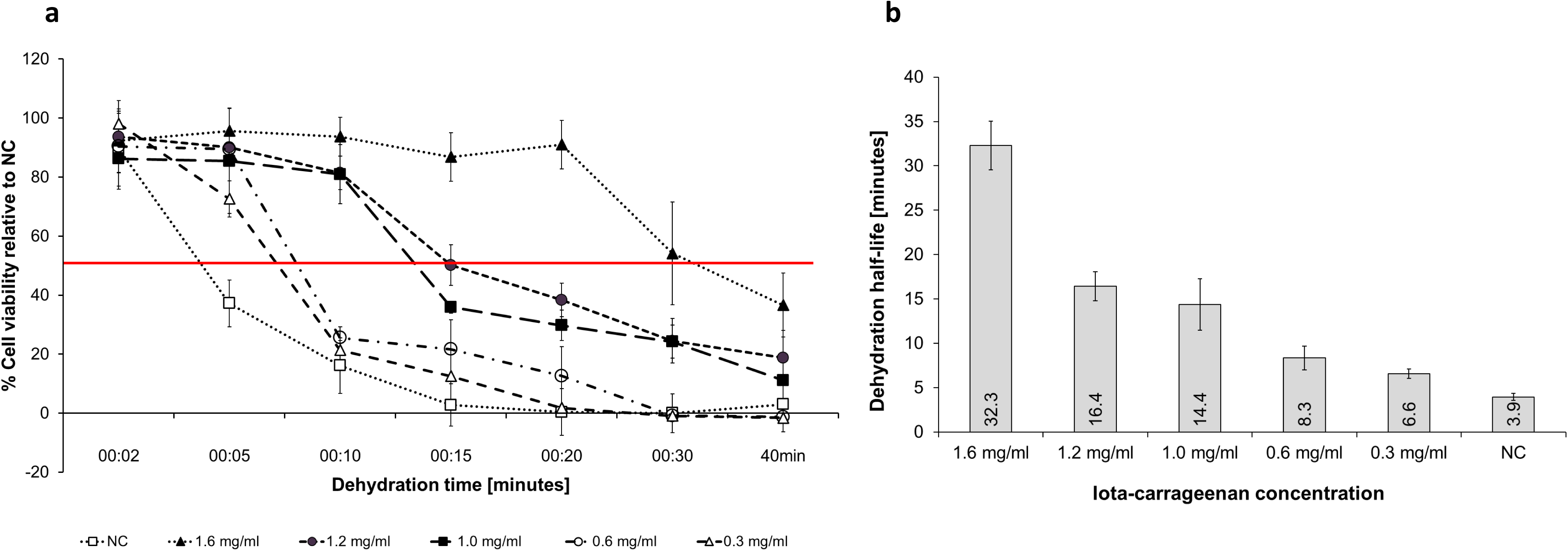
Iota-carrageenan protects corneal cells from dehydration in a concentration-dependant manner. IM-HCEpiC were pre-incubated with indicated iota-carrageenan concentrations in cell culture medium for 2h and then dehydrated for 2-40 min by removing the medium and placing the culture plates without lid in a dry incubator. **a:** Viability was assessed photometrically using a resazurin based assay after dehydrating cells for indicated time periods. Viability is given relative to the viability of the respective sample positive control (i.e., cells pre-treated with same iota-carrageenan concentration as respective sample but did not undergo dehydration). Error bars represent the standard deviation across all replicates (n = 6-12). The horizontal red line represent 50% cell viability. **b:** Survival half-life defined as time after which 50% of cells were still metabolic active after pretreatment with the tested concentrations of iota-carrageenan. Error bars represent 95% CI. NC = negative control, cells pre-treated with cell culture medium w/o iota-carrageenan, followed by dehydration.

As shown in **figure 3**, dehydration caused cells to shrink and subsequently die, leading to reduced cell density, when they had been pre-treated with medium only, while pre-incubation with 1.6 mg/ml iota-carrageenan prevents these changes and maintains cell density and healthy cell morphology.

**Figure 3:**
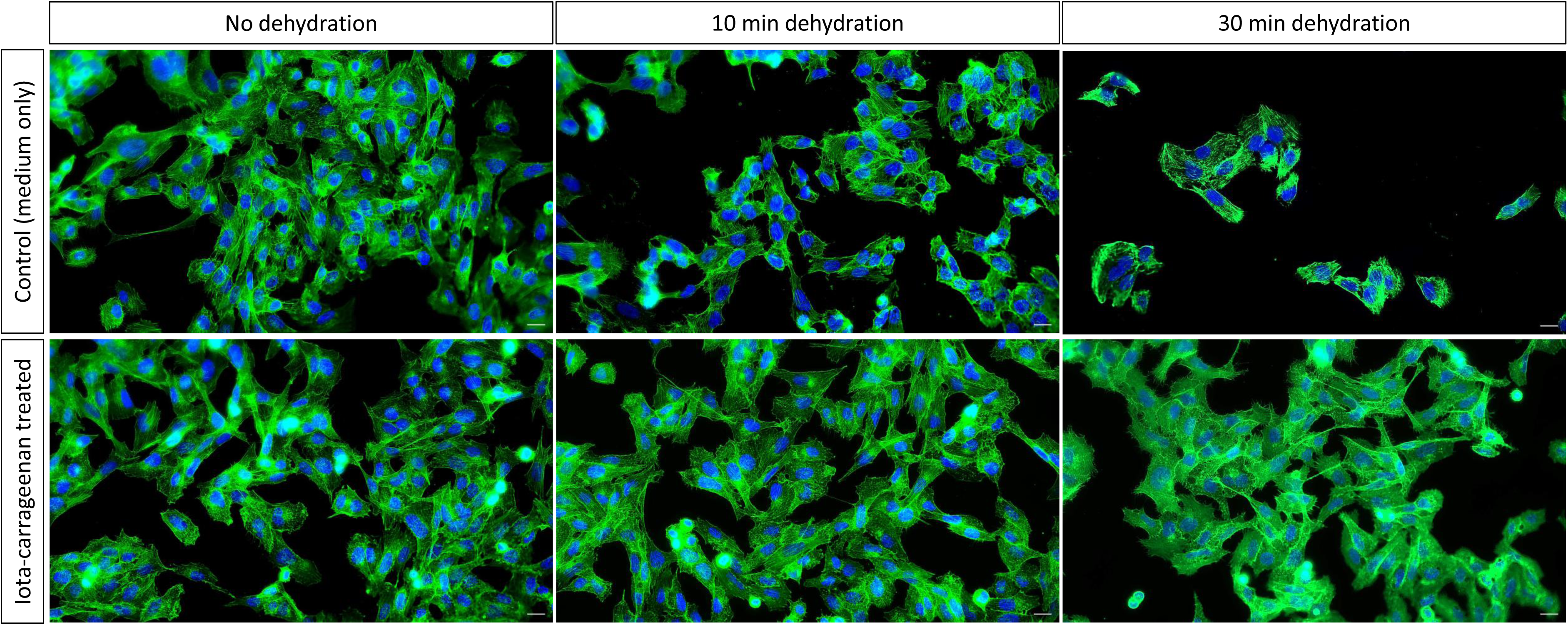
Pre-treatment with iota-carrageenan eye drops protects corneal cells from dehydration-induced morphological changes. IM-HCEpiC were pre-incubated with 3.2 mg/ml iota-carrageenan mixed 1:1 with cell culture medium (giving a final iota-carrageenan concentration of 1.6 mg/ml) for 2h and then dehydrated for 10 or 30 min. Actin fibers were stained with phalloidin (green) and nuclei with DAPI (blue) to visualize morphology. Scale bare is 20 µm.

Next, we applied a porcine *ex vivo* model to validate the results obtained *in vitro*. **Figure 4a-c** illustrates the experimental set-up. Eyes were extracted from the skulls of freshly slaughtered swine. After transport to the laboratory, eyes were visually inspected for ocular surface damage, and only undamaged eyes were used. Eye lids were pinned open and eyes were dehydrated by exposure to laminar air flow for 6 h. During this period, eyes were treated in 15 min intervals with 1 drop of either 0.9% NaCl or of 0.6, 2.0 or 3.2 mg/ml iota-carrageenan dissolved in isotonic buffer. In contrast to the *in vitro* cell culture experiment where iota-carrageenan was dissolved in AD and 1:1 mixed with cell culture medium to maintain cell viability, for *ex vivo* application iota-carrageenan was dissolved in an isotonic buffered solution compatible for ophthalmic application, i.e., with neutral pH containing sorbitol, di-sodium hydrogen phosphate dihydrate, citric acid monohydrate, and sodium- ethylenediaminetetraacetic acid (EDTA) in AD. After 6 hours of dehydration, eyes were stained with fluorescein to assess and quantify corneal damage caused by the desiccation. Fluorescein penetrates poorly into the lipid layer of the corneal epithelium, and therefore, does not stain intact cornea, but can only penetrate and stain damaged cornea where cell-to-cell junctions are disrupted (Srinivas and Rao, 2023). As shown in **figure 5a**, no obvious cornea damage was seen before dehydration. After 6 h dehydration, a very small extent of damage was observed in eyes repeatedly treated with iota-carrageenan eye drops during dehydration. In contrast, the control eyes treated with 0.9% NaCl had acquired widespread corneal damage, as indicated by white arrows. The extent of damage can be quantified by dye extraction from cornea punches after fluorescein staining. **Figure 5b** shows the amount of fluorescein extracted from cornea punches, expressed as percentage of NaCl treated control eyes. The amount of dye that penetrates the corneal epithelium and hence can be extracted from cornea samples is proportional to the corneal damage acquired during dehydration. Repeated treatment with iota-carrageenan protected porcine eyes from dehydration-induced cornea damage. With all three tested concentrations of iota-carrageenan, the amount of extracted dye, i.e., of corneal damage, was significantly reduced compared to control eyes treated with physiologic saline solution. As shown before *in vitro*, there was a clear trend towards dose dependence of the effect, even though the difference in effect size between concentrations was just short of being statistically significant (p value for comparison between 0.6 and 3.2 mg/ml = 0.059) based on the available number of biologic replicates. The baseline value has been derived from eyes not subjected to dehydration and demonstrates the effectiveness of the dehydration treatment for inflicting corneal damage.

**Figure 4:**
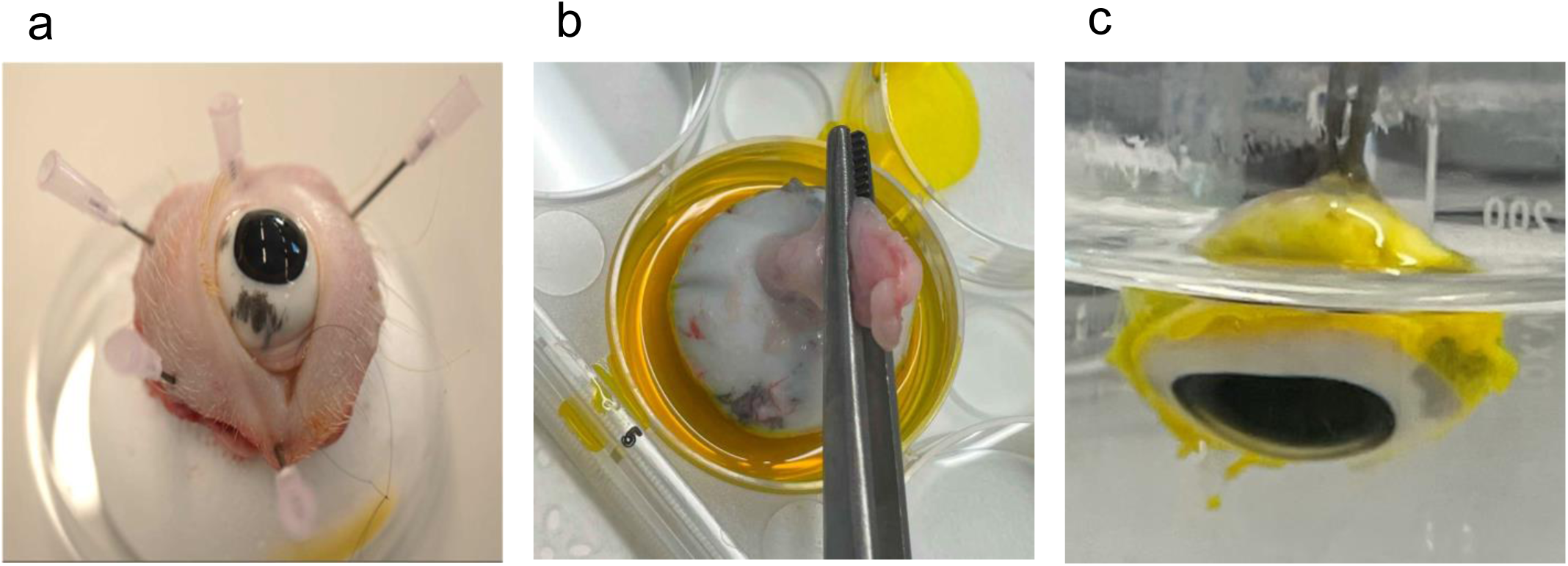
Experimental set-up of induction and detection of ocular surface damage in a porcine *ex vivo* model. Eyes extracted from freshly slaughtered swine were dehydrated by exposure to laminar air flow for 6 h under repeated instillation of 0.9% NaCl or iota-carrageenan in isotonic buffer. Eyes were visually inspected for ocular surface damage before and after dehydration. Corneal damage was quantified by dye extraction from cornea punches after fluorescein staining of eyeballs. **a-c** Handling of porcine eyes. **a:** Porcine eye were pinned open and exposed to air for 6 h. Every 15 min, 1 drop of 0.9% NaCl or iota-carrageenan solution was instilled into the eye and evenly spread by closing and opening the nictitating membrane. **b:** Immersion of eyeballs in 0.1% fluorescein solution. **c**: Washing of fluorescein-stained eyeballs. After fluorescein staining, corneas were dissected, circular punches were taken, and fluorescein was extracted from punches and quantified.

**Figure 5:**
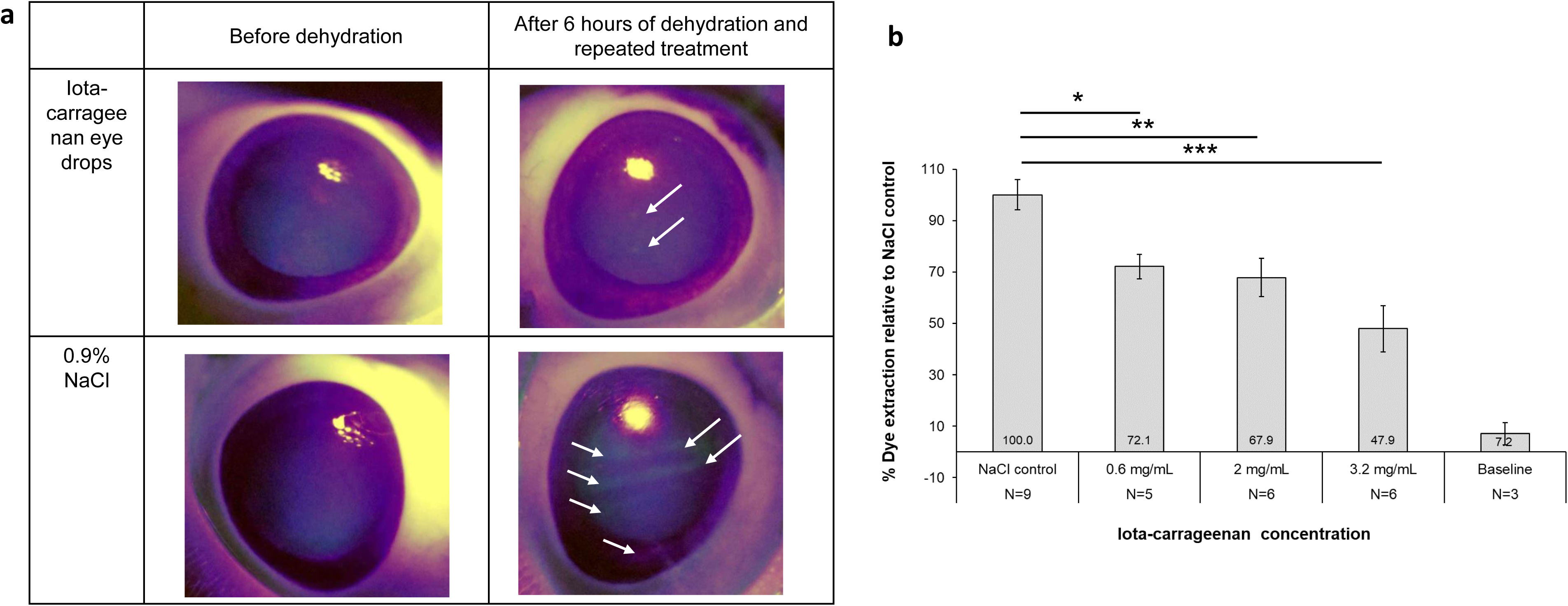
Iota-carrageenan provides dose-dependent protection of the cornea from dehydration-induced ocular surface damage in a porcine ex vivo ocular dehydration model. **a:** Freshly extracted porcine eyes were examined for cornea damage under a black light torch after instilling 25 µl fluorescein solution (left column). Only undamaged eyes (Oxford scale grading 0 or 1) were used in the experiment. After 6 hours of dehydration under repeated instillation of 0.9% NaCl or 3.2 mg/ml iota-carrageenan eye drops, eyes were again visually inspected under black light after instilling of 25 µl fluorescein solution (right column). Widespread corneal damage is seen after dehydration in NaCl-treated eyes, but not in eyes treated with 3.2 mg/ml iota-carrageenan eye drops (white arrows indicate corneal damage). **b:** The amount of fluorescein extracted from cornea punch samples after dehydration and immersion into a 0.1% fluorescein solution, expressed as percentage of NaCl treated controls, is proportional to the corneal damage acquired during dehydration. Error bars represent the SEM. N=number of eyes used per treatment group. * p ≤ 0.05; ** p ≤ 0.01;*** p ≤ 0.001. Baseline value (=before dehydration) is significantly different from all other samples.

In sum, our data demonstrate the capacity of iota-carrageenan to protect against dehydration in an *in vitro* cell culture system of human cornea cells as well as in an *ex vivo* animal model. Based on these *in vitro* and *ex vivo* results, 3.2 mg/ml was determined to be the most effective concentration of iota-carrageenan to protect ocular cells and tissue from dehydration-induced damage.

## 5. Discussion

To restore and maintain a healthy ocular surface, it is essential that both the cornea and conjunctiva remain continuously hydrated - a function normally provided by the tear film. Recent consensus identifies tear film instability as the core mechanism underlying DED. This understanding has led to the development of tear film-oriented diagnosis, which seeks to clarify the specific causes of DED in individual patients by focusing on the distinct layers of the tear film. Consequently, tear-film-oriented DED therapy focuses on restoring tear film homeostasis and ensuring adequate lubrication of the ocular surface (Dash and Choudhury, 2024; Safir et al., 2024; Shimazaki, 2018).

The research presented in this article investigated the protective effect of iota-carrageenan against air-induced desiccation and evaluates its potential as an ocular lubricant for the relief of mild to moderate DED.

Our data show that in a cell culture model of corneal epithelial cells, pre-treatment of cultured cells with an iota-carrageenan solution dramatically enhanced cell survival after dehydration. The magnitude of the protective effect was comparable to high molecular weight hyaluronic acid, and was superior to all other tested natural polymers, including other widely used ingredients of eye drops used for DED treatments. Moreover, our results highlighted significant differences in the effectiveness of hyaluronic acid based on its molecular weight. The protective effect of iota-carrageenan was dose-dependent, as higher concentrations correlated with increased survival rates. Moreover, iota-carrageenan significantly inhibited the desiccation-stress induced accumulation of corneal damage in an *ex vivo* porcine eye model also in a dose dependent manner. In sum, the data presented here suggest that iota-carrageenan has a moisturizing and protective effect that prevents cornea cells from dehydration and, in consequence, from dehydration-induced cell death and tissue damage. Of note, the highest concentration used in the *ex vivo* experiment (3.2 mg/ml; maximum possible concentration for technical reasons) exceeds the highest concentration in the cell culture experiment described above (1.6 mg/ml). This was intended to compensate for the dilution with the lacrimal fluid and the rapid restoration of the tear film that washes away a considerable proportion of the administered solution (Gaudana et al., 2010).

The texture properties of carrageenans have long been known and exploited in the food and the cosmetics industry. Carrageenans are known to retain moisture, enhance viscosity combined with shear thinning behavior, and have mucoadhesive properties that are also exploited in various forms of drug delivery (Bonferoni et al., 2004; Volod’ko et al., 2024). We hypothesize that these properties together contribute to the effect observed in our experiments: Carrageenan forms a mucoadhesive in situ film on the cellular monolayer and on the *ex vivo* porcine eye, it enhances viscosity of the solution, improves surface retention, and counteracts dehydration due to water-retention. We claim that these mechanisms should work equally well in the human eye, however this shall be subject to further studies.

Hyaluronic acid containing eye drops have become the first choice for tear substitution in many countries. According to literature, the average molecular weight of hyaluronic acid determines the therapeutic efficacy of such eye drops (Salzillo et al., 2016), and high molecular weight hyaluronic acid has enhanced mucoadhesion and moisturizing capacity compared to the low molecular weight form (Guarise et al., 2023). Hyaluronic acid is a viscosity enhancer used to form gel-like ophthalmic solutions which may cause, especially with high molecular weight, blurred vision and a foreign body sensation, especially immediately after instillation (Müller-Lierheim, 2020). In contrast, in situ gelling systems are generally considered more comfortable because they are installed as low-viscosity liquids causing minimized immediate vision disruption (Cassano et al., 2021). In our experiments, the viscosity of the high molecular weight hyaluronic acid solution was 9 mPas and that of the iota-carrageenan solution 2 mPas, measured at 33°C and a shear rate of 10 1/s with an Rotavisc lo-vi (IKA) with ELVAS-1 extension (data not shown). We have shown *in vitro* that iota-carrageenan is equal to high molecular weight (1.2-1.5 MDa) hyaluronic acid in terms of cytoprotection under dehydration at a concentration of 1.2 mg/ml and therefore expect similar therapeutic efficacy of iota-carrageenan-based eye drops as reported for hyaluronic acid-based eye drops. It is important to note that our *ex vivo* model represents only a simplified approximation of the eye *in vivo*, where continuous secretion from lacrimal and Meibomian glands as well as regular blinking occur. To partially compensate for the lack of natural blinking, our experimental set-up included mechanic blinking with the lid to facilitate the distribution of test solutions. No tears replenishment was performed in the *ex vivo* eyes which on one hand prevented wash out of the treatment but also led to rapid cornea dehydration. We tried to compensate for this with repeated installations of the respective test solutions. Nevertheless, this set up does not sufficiently consider the natural tear clearance and the washout processes that occur *in vivo* .

Further studies are required to shed light on the safety, tolerability and therapeutic effectiveness of iota-carrageenan eye drops in the treatment of DED in humans.

## 6. Conclusions

Iota-carrageenan exhibits significant dose-dependent protection against desiccation-induced cell death and damage in both in vitro cultured human corneal cells and an ex vivo porcine eye model. Iota-carrageenan closely mimics natural tear fluid due to its mucoadhesive properties, enabling it to substitute lost tear fluid. By lubricating and hydrating the cornea and conjunctiva, it supports a healthy ocular surface and additionally protects the eye surface from dehydration-induced damage.

Our findings position iota-carrageenan as a promising novel ingredient for DED-targeted eye drops. Its dual capacity to replace tear fluid and shield the ocular surface from dehydration underscores its potential as an advanced therapeutic agent.

## 7. Declarations

### 7.1 Acknowledgement

The authors thank Dorian Winter, Nicole Reiterer-Felber and Fabian Wallisch for excellent technical and scientific support.

### 7.2 Statement of Ethics

Regulations on the protection of animals used for scientific purposes do not apply to the present study, since tissues of animal origin used in this study were derived from the local slaughterhouse as waste product from food production under adherence to national laws on the protection of animals in connection with slaughtering.

### 7.5 Data Availability Statement

Data related to this manuscript can be made available from the corresponding author upon reasonable request.

### 7.6 Author Contributions

SRN: conceptualization, project administration, supervision, data analysis, writing – review and editing; AR: methodology, supervision, data analysis, writing – review and editing; CK: methodology, data analysis, writing – review and editing; AD: methodology, data analysis, writing – review and editing; MKS: methodology, data analysis, writing – review and editing, HD: data analysis, visualization, writing – original draft, writing – review and editing; EPG: conceptualization, supervision, data analysis, writing – review and editing.

